# The dentate gyrus provides flexibility for efficient spatial navigation

**DOI:** 10.1101/2024.12.31.630892

**Authors:** Natalia Soldi, Sabrina Benas, Emilio Kropff, Alejandro F. Schinder, Verónica C. Piatti

## Abstract

The hippocampus plays a critical role in spatial navigation and declarative memory. The dentate gyrus is the neurogenic region of the hippocampal formation and it has long been implicated in the fine separation of similar contexts or close object locations. However, it is unclear how an accurate discrimination could be beneficial to a goal-guided behavior in a changing environment. Therefore, we used chemogenetic inhibition to study the role of the dentate gyrus in a goal-guided spatial navigation paradigm over a familiar but dynamic crossword maze. Mice were challenged to localize a novel reward location from alternative pathways in two versions of the task with particular configurations in each experimental day. In the simple task, the two optimal paths to the goal shared some segments in their trajectory. In a more complex task, optimal trajectories demanded completely different directions to the reward location. Overall, mice with chemogenetic inhibition of the dentate gyrus were able to learn all the routes regardless the complexity of the task, similarly to control animals. However, after having solved a first route in the complex task, mice with dentate gyrus inhibition displayed an impairment to efficiently navigate over the alternate path. Our results demonstrate a role of the dentate gyrus in cognitive flexibility required to reach a goal in a changing familiar environment.

## INTRODUCTION

The dentate gyrus is a sparse competitive network suitable for input discrimination, which results in what is known as pattern separation (GoodSmith et al., 2017; Piatti, Ewell, & Leutgeb, 2013). In computational terms, this algorithm points to the notion that similar stimuli are orthogonalized into completely dissimilar outputs (Cayco-Gajic & Silver, 2019; Treves & Rolls, 1994). In regard to behavior, this concept has been translated as the capacity to discriminate similar contexts or close object locations (Kesner, 2013; Sahay, Wilson, & Hen, 2011; Yassa & Stark, 2011). Rodents with manipulations that impair the dentate gyrus cannot discriminate subtle object displacements or reconfiguration of the same objects in their familiar environment (Bekinschtein et al., 2013; Kesner, Taylor, Hoge, & Andy, 2015). Although these metric differences could be as small as 2 cm, they are novel and because they share some features of past experiences in the same environment they are “common novelties” (Duszkiewicz, McNamara, Takeuchi, & Genzel, 2019; Yamasaki & Takeuchi, 2017). In a continuously changing environment, the behavioral discrimination of subtle common novelties could become a disadvantage for survival. In fact, it has been observed that the opposite behavior is advantageous, called generalization, which is the ability to extend memories from similar past events to solve new situations in the future. The imbalance of these opposite processes has been reported in anxiety disorders and psychosis, such as schizophrenia. In anxiety, there is overgeneralization, while in schizophrenia there is strong discrimination (Ivleva et al., 2012; Kheirbek, Klemenhagen, Sahay, & Hen, 2012). Intriguingly, discrimination and generalization are functions of different neuronal circuits of the dentate gyrus (Inoue & Watanabe, 2014; Sun et al., 2020). Therefore, what is the dentate gyrus role in a changing environment where animals must survive based on their choices?

In cognitive tasks in which the goal is to avoid an aversive stimulus or find a reward, the dentate gyrus is crucial to discriminate as well as to generalize common novelties to reach the purpose. Thus, it is critical to discriminate between an aversive context and a new similar one that is not aversive (McHugh et al., 2007). Likewise, it is necessary to distinguish a rewarded location from an adjacent similar place (Gilbert, Kesner, & Lee, 2001; A. M. Morris, Churchwell, Kesner, & Gilbert, 2012). The dentate gyrus is also necessary to generalize when the reward location changed keeping the same cues configuration around it (Inoue & Watanabe, 2014). In summary, this hippocampal region has been shown to have versatile functions accordingly to the animal needs for survival, which implies an adaptive role of this region. All these studies have in common that rodents were trained in the task for several expositions in which mnemonic processing of several stimuli could have been taken place beyond an algorithm of input orthogonalization.

Adaptation to an ever-changing world is critical for survival, and this ability requires the continuous update and selection of memories (R. G. Morris, 2006). Understanding how memories guide behavior in novel situations of the daily life will provide insights into a fundamental aspect of adaptive animal behavior. The hippocampus has been fundamental for these cognitive processes along evolution (Treves, Tashiro, Witter, & Moser, 2008). For instance, mice are impaired to learn a new reward position in a familiar crossword maze when the CA1 region of the hippocampus is optically silenced (McNamara, Tejero-Cantero, Trouche, Campo-Urriza, & Dupret, 2014). However, whether other hippocampal areas are needed to solve this task remains unclear. Here, we investigated the requirement of the dentate gyrus in the same challenge and a new version of the task, using chemogenetic inhibition mediated by the synthetic receptor hM4Di (Urban & Roth, 2015). Interestingly, dentate gyrus shutdown did not alter overall learning in the behavioral task. Though, the navigation efficiency towards the goal was impaired when mice had to reach it via an entirely different route. These results support the notion that the dentate gyrus provides cognitive flexibility bearing an adaptive value for navigation in a changing environment.

## MATERIAL AND METHODS

### Animals

All animal experiments were performed following the approved guidelines of the Institutional Animal Care and Use Committee of the Leloir Institute (CICUAL-FIL 98), according to the Principles for Biomedical Research involving animals of the Council for International Organizations for Medical Sciences and provisions stated in the *Guide for the Care and Use of Laboratory Animals*. Leloir Institute is approved as a foreign facility by the Office of Laboratory Animal Welfare of the US National Institutes of Health (F18-00411).

The C57BL/6J mouse line from Jackson Laboratory used in this study was bred in our animal facility for over ten generations. Mice were socially housed on an inverted 12 h dark/12 h light cycle room (lights on 08:00 p.m.) from their weaning until their sixth week old. At that age they were isolated and hosted with an individual running wheel until the end of the experiments. Water and food were provided ad libitum and only restricted during the behavioral training between a range of 85 - 95 % of their body weight. Behavior was conducted using male (10) and female (11) mice aged 8-12 weeks during their dark phase. From the weaning day until the last day of each mouse, they were kindly treated in their care for the Vivarium technicians to get used to a good human connection.

### Viruses

Adeno-associated viral vectors (AAV) (8)-CaMKIIa-hM4D(Gi)-mCherry (Catalog # 50477-AAV8) and AAV (8)-hSyn-mCherry (Catalog #114472-AAV8) were purchased from Addgene. AAV (9)-pCAG-FLEX-tdTomato-WPRE were a gift from Silvia Arber laboratory based on a backbone vector from Allen Brain (AAV.CAG.FLEX.tdTomato.WPRE.bGH) (JA Harris, SW Oh, H Zeng, Curr. Protoc. Neurosci., 2012). AAV (9)-hSyn-double floxed HA-hM4Di-WPR was a gift from Silvia Arber laboratory. The retrovirus (rv)-CAG-Cre was made in the laboratory with the same protocol already published (Piatti et al., 2011).

### Stereotaxic surgery

Mice were anesthetized using ketamine-xylazine (150 mg ketamine/15 mg xylazine in 10 mL saline/g) and then placed into a stereotaxic frame (Stoelting, Catalog # 51730D). Eye drops were applied to prevent drying. From Bregma the coordinates used for the bilateral viruses deliver into both dentate gyrus were −2 mm anteroposterior, ± 1.5 mm mediolateral, and −1.9 mm dorsoventral according to the mouse brain atlas. The viral infusion was of 0.5μl - 1.5μl at 0.1 μl/min in each site using sterile-calibrated microcapillary pipettes through stereotaxic references accordingly to the experimental group. The needle remained in place for an additional 2 min before being slowly withdrawn. The skull was kept moist throughout the operation, then sutured, and mice were transferred back to their home cages. The surgeries were all performed when mice were 4 weeks old and the behavioral training started 6 weeks after viral injection (Fig. 1A). The early surgery time and the extended timing between the viral infections and the behavior experiments onset were to provide extra time for recovery since the reported publication that some AAVs ablated adult neurogenesis (Johnston et al., 2021).

**Figure 1:**
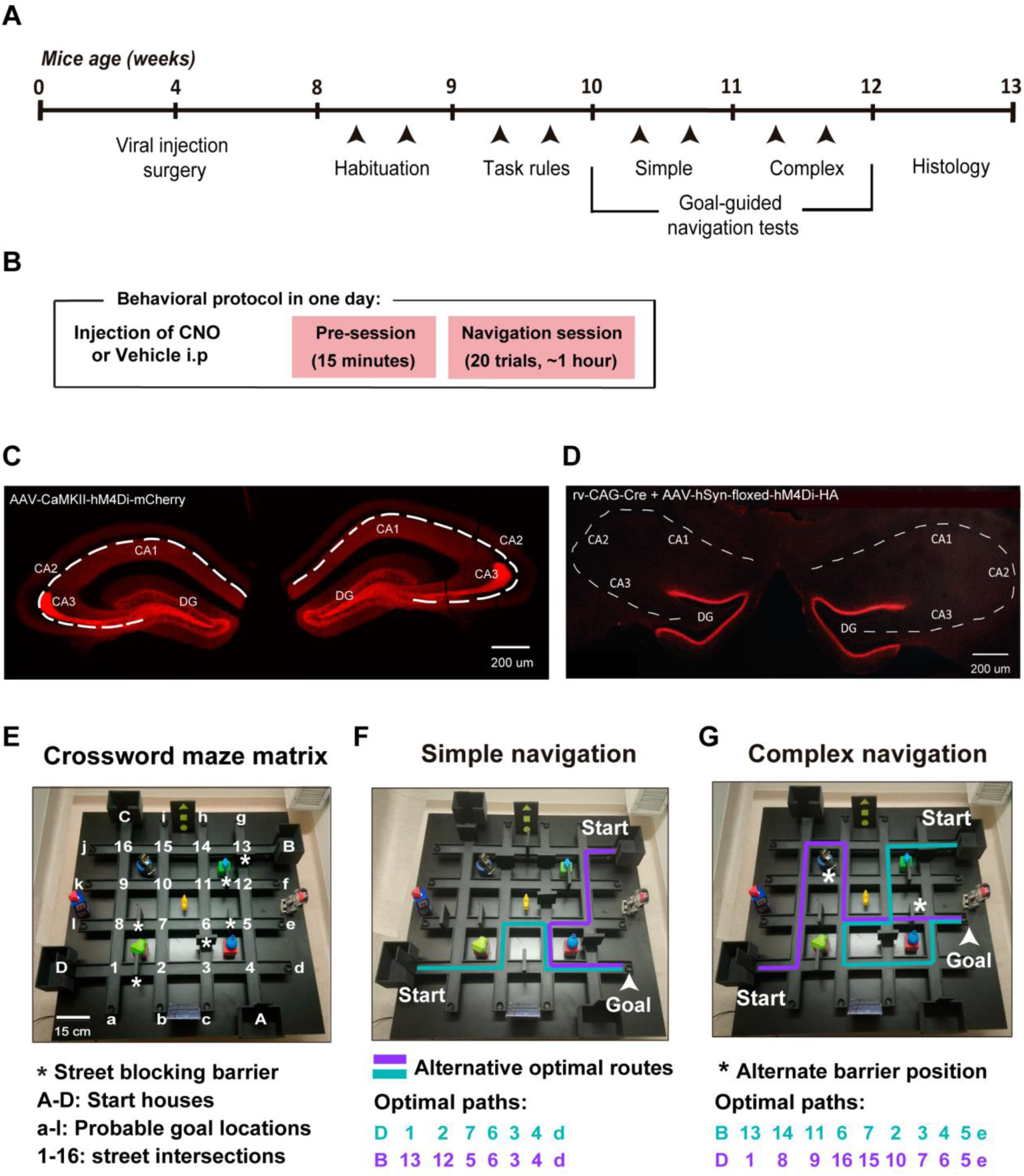
Goal-guided navigation tasks in the crossword maze and experimental strategy. **A**. Behavioral timeline of experiments in all C57Bl6/j mice used in this study. Behavioral experiments were done when mice were 10th - 12th weeks old. Black arrowheads point to one behavioral experimental day in the maze. **B.** All mice did both tests twice, one injected with a vehicle and another with CNO randomly chosen. These injections were previous to the spatial navigation sessions. In each experimental day, the same protocol was done: the maze was rotated, a different set of barriers was used to generate new optimal routes towards a unique new reward location (common novelty in this paradigm) fixed during one session. **C.** Coronal hippocampal section of a representative mouse injected bilaterally with the adenovirus AAV-CaMKII-hM4Di-mCherry. Note the hM4Di-mCherry expression in the dentate gyrus dendritic field, mossy fibers, commissural axons and the Schaffer collateral of CA3. **D.** HA-immunohistochemical section of an example bilateral coronal hippocampus of a mouse injected with a viral mix of a RV-CAG-Cre plus an AAV-hSyn-double floxed-hM4Di-HA. Note the specific hM4Di-HA expression in the dentate gyrus commissural fibers. **E.** Photo of the crossword maze with the matrix intersection layout. The street intersections are “1 to 16”, the alley ends are “a to l” and the starting points are “A to D”. **F.** Photo of the maze with a simple navigation task configuration layout. Example of one pair of optimal routes drawn from the two available houses in one session. The route color means the alternative pathways that were randomly assigned for each day. White arrowhead points to the reward position. At the bottom the optimal sequence of intersections from the starting house to the goal are depicted as examples from that particular session. **G.** Example of one complex navigation layout with the pair optimal routes and the reward location coded as F. Note that in this case there are 5 fixed blocking barriers and 1 mobile barrier that alternate between two positions (*) depending on the alternating trial. The sequences for the optimal routes of this configuration are written at the bottom.

### Experimental groups

There were 4 groups of mice accordingly to the type and anatomic expression of the fluorescent markers injected with the viral vectors. Two experimental groups and two controls. The experimental groups were infected with viral vectors to drive the expression of inhibitory hM4Di receptor in the neuronal populations of the dentate gyrus. One experimental group consisting of 10 mice injected with AAV(8)-CaMKIIa-hM4D(Gi)-mCherry (0.5μl at 0.1 μl/min, diluted in 1:10 in PBS) this infection has a broad expression in the dentate gyrus and some CA3 neurons (Fig. 1C). Therefore, another experimental group was done to restrict the hM4D only to dentate neurons. This group were 5 mice which were injected with a mixture of viral vectors (1.5 μl at 0.15 μl/min). The combination was 2/3 of the final volume of the rv-CAG-Cre plus 1/3 of the AAV (9)-hSyn-double floxed HA-hM4Di-WPR. Retrovirus are known to infect cells in division but this particular preparation was unspecific for all neurons (Fig. S1 A, B). This combination resulted in a localized hM4D expression in the commissural fibers of the dentate gyrus (Fig. 1D). The control groups were mice also infected with AAVs but driving the expression of fluorescent markers without the inhibitory receptor. Controls were 6 mice total that were infected with: AAV (8)-hSyn-mCherry (0.5μl at 0.1 μl/min) (N = 4 animals) and a mixture of 2/3 of rv-CAG-Cre plus 1/3 of AAV (9)-pCAG-FLEX-tdTomato-WPRE diluted in 1:3 in PBS (1.5 μl at 0.15 μl/min) (N = 2 mice). All these 6 mice have a similar pattern of broad expression of fluorescent markers into dentate gyrus and CA3 cells, therefore they were pooled together to a unique control group of the hM4D ligand (Fig. S1 C).

### Chemogenetic manipulation

The strategy used in this study to modulate the neuronal activity of the dentate gyrus was the chemogenetic inhibition by the synthetic receptor hM4Di (Urban & Roth, 2015). AAVs and retrovirus were injected at early age to carry the receptor into both dentate gyrus of wild type mice. Behavioral tests were performed at least 6 weeks after viral infection. At the experimental day mice were injected intraperitoneal (i.p.) with 5 mg/kg clozapine N-oxide (CNO) or vehicle solution (saline) chosen randomly. The time of injection in a single experimental day was 120 min before the first trial of the navigation session. CNO solution was 5 mg/ml of CNO (Enzo, Catalog # BML-NS105-0025) in saline solution and it was prepared fresh for each experiment with a maximum of use 10 days.

### Behavioral assays

In the 7^th^ week of life, mice were gentled handled by the experimenter for at least 2 days (10 min) in their home cage in the inverted-light cycle room under red light illumination. During this timing, mice were provided with pellets embedded in chocolate milk or sunflower seeds to assess the preferred food reward for each animal. At 8 weeks old, the habituation was done in 2 days (30 min each day) (Fig. 1 A). In the next week, mice were familiarized with the paradigm in the maze for another 2 days (∼1 hr each training) where they learn the task rules. Finally, the simple and complex goal-guided navigation tests were done between their 10^th^ and 12^th^ weeks old (each week, 2 days per test). All behavioral test were done 120 minutes after the saline or CNO i.p. injection (randomized). This i.p. injection was performed in their accommodation room by another person different than the experimenter to avoid a stressful relation with the animal (Fig. 1 B). Every animal manipulation was done in their dark phase, generally between 14 – 20 hrs. Animals were transported between rooms in a dark bucket. The behavioral room had a dim light (simile moon light) and a camara on the roof to film the animal behavior on the maze.

#### Crossword maze

The crossword-like maze used in this study was an adapted version of the crossword maze previously used (McNamara et al., 2014). It consisted of a symmetrical square of 1.2 x 1.2 meters made of 4 equidistant linear tracks orthogonalized with another 4, leading to 16 perpendicular intersections between tracks. In each corner of the square, at the end of their lateral tracks, there were 4 square houses with manually modulable doors. The rest of the tracks finished in a free alley with a black little cup at their end, generating 12 possible positions for the reward. Intersections, streets ends and houses were labeled with numbers and letters turning the maze into a matrix in order to track the animal path with a sequence of numbers and letters (Fig. 1 E). Each house had 15 × 10 cm base, with 15 cm high walls. The width of each track was 5 cm with a 1.5-cm-high rim along the edges. Mobile barriers of 10 × 10 cm square and 2 cm width were available to insert on the streets in order to block the path, allowing to make different configuration of routes for goal-guided navigation. The maze had local clues spread between the different street block and distal clues placed in the walls of the room together with equipment and the experimenter in one side of the maze. To promote allocentric spatial navigation, the maze was rotated in order to change the relative positions between the distal cues and the local ones, at the beginning of each behavioral day.

#### Habituation

Each mouse was taken to the behavioral room once per day during 2 days of the week. The habituation had 4 stages. For the first 10 minutes, the animal was gently transferred to the rest tray in a pedestal. Then it was manually transferred to one of the starting houses and it was allowed to explore the maze without any door and barrier. Black little cups were positioned at the 12 alley ends carrying the chocolate milk or a piece of a sunflower seed in all of them accordingly to the mouse preference. After 10 minutes of this first exploration, the mouse was transferred to the rest tray for one minute while the maze was clean and the cups were all filled again with the corresponding reward. Subsequently, another 10 minutes of free exploration was allowed on the maze in the same way; without barriers or doors. In this case, the animal was positioned in the house that was opposite to the previous one chosen for the first exploration (Fig 1 E, houses combination was A - C or B - D). Lastly, the habituation session ended with the mouse in the rest tray for 2 minutes and then they were taken back to their home cages in their accommodation room.

#### Task rules

The familiarization consisted on 2 behavioral sessions separated in two different days. The animals performed a goal-guided spatial behavior, the simple task (see below, “Goal-guided spatial behavior”), but in this case mice were taught to find the reward in one of the cups and learn the task rules. The particularity of the familiarization session was that the animal could be guided by the experimenter towards the reward. This happened in the cases that the mouse performed 3 consecutive trials without finding the rewarded cup, in order to avoid frustration. In addition, in these sessions the animals were taught not to climb the barriers, the house doors and walls and not to jump out of the maze or over the local clues (toys). If one of these rules was broken, the experimenter picked the mouse out of the maze and pressed its tail-tip as a punishment. That trial was stopped and considered “Out” and the mouse return to the rest tray until the next trial started. For these sessions and so on, animals were food restricted (see section “Animals”) from 2 days in advance.

#### Goal-guided spatial behavior

The simple and complex navigation tasks consisted in two sessions of 20 trials each that were done in different days, one day corresponding to the vehicle injection and the other to CNO (control conditions or dentate gyrus inhibition, respectively). The crossword maze was configured to generate a new pair of optimal routes to reach a unique reward position from opposite houses in each session of the simple as well the complex task. The configurations were achieved due to the mobile barriers that were disposed in specific arrangements in order to create different routes from opposite houses to a new goal location in each session. The goal place was constant in the same track-tip cup along the hole session and the others 11 cups were present but empty. The behavioral test started with the mouse spending 10 min rest in the pedestal, continued by a 15 min of pre-navigation of free exploration in the maze (Fig. 1 B). The animal was positioned in the center of the maze, which was without barriers and cups and all the houses were closed. Then it was returned to the rest tray for 5 min while the maze was cleaned with alcohol 70 % and the barriers, cups and the reward were set up. In addition, a little plastic box with perforations containing a small amount of the reward without having it accessible for the animals was used to signal the onset of the trial to them. This sign box was introduced in one of the two opposite houses randomly chosen. The navigation session started placing the mouse in this house with the sign box and closing its door. After 10 s the sign box was removed and the door was manually raised, allowing the animal to explore the maze. The remaining three houses stayed closed for the whole trial. The mouse had 180 s (maximum trial time) to find the goal at the novel track tip. At the next trial, the mouse was placed inside the second opposite house selected for that day with the sign box and performed the same procedure as before. The alternation between houses continued until 20 trials were done (10 trials for each house, 1/1.5 hour approximately, Fig. 1 B). Finally, the mouse spent 60 min in the rest tray before it went back to its home cage.

Each trial could finish being a correct or successful trial when the mouse achieved its reward before the 180 s. In the case the animal did not reach the goal in the maximum time was a time out trial, it was removed from the maze to the rest tray in the pedestal. A third situation of trial ending could happen when the animal violated one of the task rules, such as climbing a barrier, so it was immediately removed to the pedestal too and it was an out trial. In the rest tray, an inter-trial-interval of 2 min was done meanwhile the experimenter filled the reward cup and cleaned the maze with alcohol 70 %. This procedure was done for most of the animal groups, except for the DG local expression of hM4D (Fig. 1 D) consisting of 5 animals, where the maze was not cleaned in between trials.

In the “**Simple navigation task”,** 10 barriers were placed in different tracks between 2 intersections and kept fixed along all the session. So, the different configurations for each session had 2 shortest routes of 6 intersections long (in average) to the reward position. As mice started from 2 opposite houses, the first direction choices towards the unique goal position were different for each path but the last 3 intersections to the goal were overlapped (Fig. 1F). There were 10 “trials out” from the total of 200 trials of the 10 analyzed mice with hM4D broad expression (Fig. 1 C) in the vehicle condition and 9 of 200 trials of the same mice in the CNO condition.

In the **“Complex navigation task”,** 6 barriers were used to generate different configurations to create 2 shortest routes of 9 intersections long (in average) to the reward position. In these configurations, 5 barriers were in fixed positions during the session and 1 barrier was mobile, alternating between two fixed positions. Each fixed position of the mobile barrier was related to one of the two starting houses and forced the animals to take a different pathway for each alternative trial from their started to goal place (Fig. 1 G). There were 8 “trials out” from the total of 300 trials of the 15 experimental mice in the vehicle condition and 11 of 300 trials of the same mice in the CNO condition.

The mice performance on the crossword maze was quantified using the 3 following parameters. The time in which the animals ended a trial, initiating from the first moment they come out of the starting house. The number of intersections traveled during one trial, obtained from the videos recorded during the session and quantified using the maze matrix (Fig. 1 E). Last, the navigation errors also scored from the same videos. An error was computed when the mouse chose a direction that took it apart from the optimal route in every intersection. An incorrect choice was considered when the animal stepped into a street with the two front feet.

### Histology and Imaging

The mice were deeply anesthetized with an i.p. injection of ketamine-xylazine and perfused intracardially with peristaltic bomb (6 ml/min) with 100 ml of heparinized (100 µl/l) saline solution, and then with 100 ml of 4% paraformaldehyde (PFA) in PBS 0.1M (pH 7.2). The brains were collected, postfixed with 4% PFA at 4℃ for an overnight, transferred to 30% sucrose in PBS until they sank to the bottom of the container, and sliced into 40 μm coronal sections using a freezing microtome (Leica, SM2000R, Germany). The sections were stored in cryoprotectant containing 30% glycerol (v/v) and 20% ethylene glycol (v/v) in PBS at - 20 ℃ until usage. Hippocampal sections from injected mice with viral vectors driving the fluorescent markers expression were washed in PBS and mounted on glass slides. After around 15 min drying, they were covered with PVA-DABCO mounting media and coverslips. The fluorescence microscope Axio Examiner.D1 (Zeiss, Germany) was used to obtain digitalized images.

Hippocampal sections from mice infected with AAV (9)-hSyn-double floxed HA-hM4Di-WPR were immunofluorescent staining for the epitope HA. The immunostaining was done on free-floating coronal sections taking a sample of 1/6 of the collected sections of the whole brain containing hippocampus in multi-well dishes. The sections were washed three times with Tris-buffered saline (TBS) (5 min each wash) and incubated with a blocking buffer containing 3% donkey serum and 0.25% Triton X-100 dissolved in TBS for 1 h. Then, they were incubated with HA primary monoclonal antibody diluted (1:250; Roche) in blocking buffer at 4 ℃ for 48 hs. After another round of blocking solution of 15 min, the sections were incubated with a fluorescent dye-conjugated secondary antibody Cy5 (1:250; Jackson ImmunoResearch) at room temperature for 2 h. Finaly, the sections were washed three times with Tris-buffered saline (TBS) (5 min each wash) and then mounted with PVA-DABCO mounting media and coverslips.). The slides were imaged using an inverted fluorescence microscope Observer (Zeiss, Jena, Germany).

### Statistics

Statistical comparisons were performed using no parametric test because there was not equal variance and independence between the groups since the same mice were control and treatment. Statistical significance was set at *p < 0.05, **p < 0.01, and ***p < 0.001. Data were presented as mean ± s.e.m. Repeated measure analysis for one or two conditions was done with Friedman test using JASP 0.16.2. Non parametric paired comparisons were performed using Wilcoxon rank-sum test and Kruskal-Wallis test with Dunn’s multiple comparison test for more than one selected comparison using GraphPad Prism 8. Comparisons were done in the cases where the standard error bars were not equal between the groups to compare and the data in the group had 5 or more samples. The shuffle analysis for the cumulative rewarded trials was performed using a Python script. Essentially, this analysis randomly shuffles the data label as “VEHICLE” and “CNO” for each mouse. This processing is performed in 1000 iterations and disrupts any systematic relationship between the treatment and the trajectory lengths, generating simulated data under the null hypothesis that there is no treatment effect. After shuffling, the same cumulative analysis is repeated on the trajectory lengths, identifying the successful trials. Finally, the difference between areas is calculated for each pair of data distribution as shown in Fig 2 D, E, F, Fig. S 1 E and compared to the observed data for statistical value. For shuffle analysis in Fig 5 B, tag-shuffles between conditions were done first, same as before. To obtain the four groups, for each iteration of shuffling we then classify in match and detour to obtain the areas of the curves as previously done.

**Figure 2:**
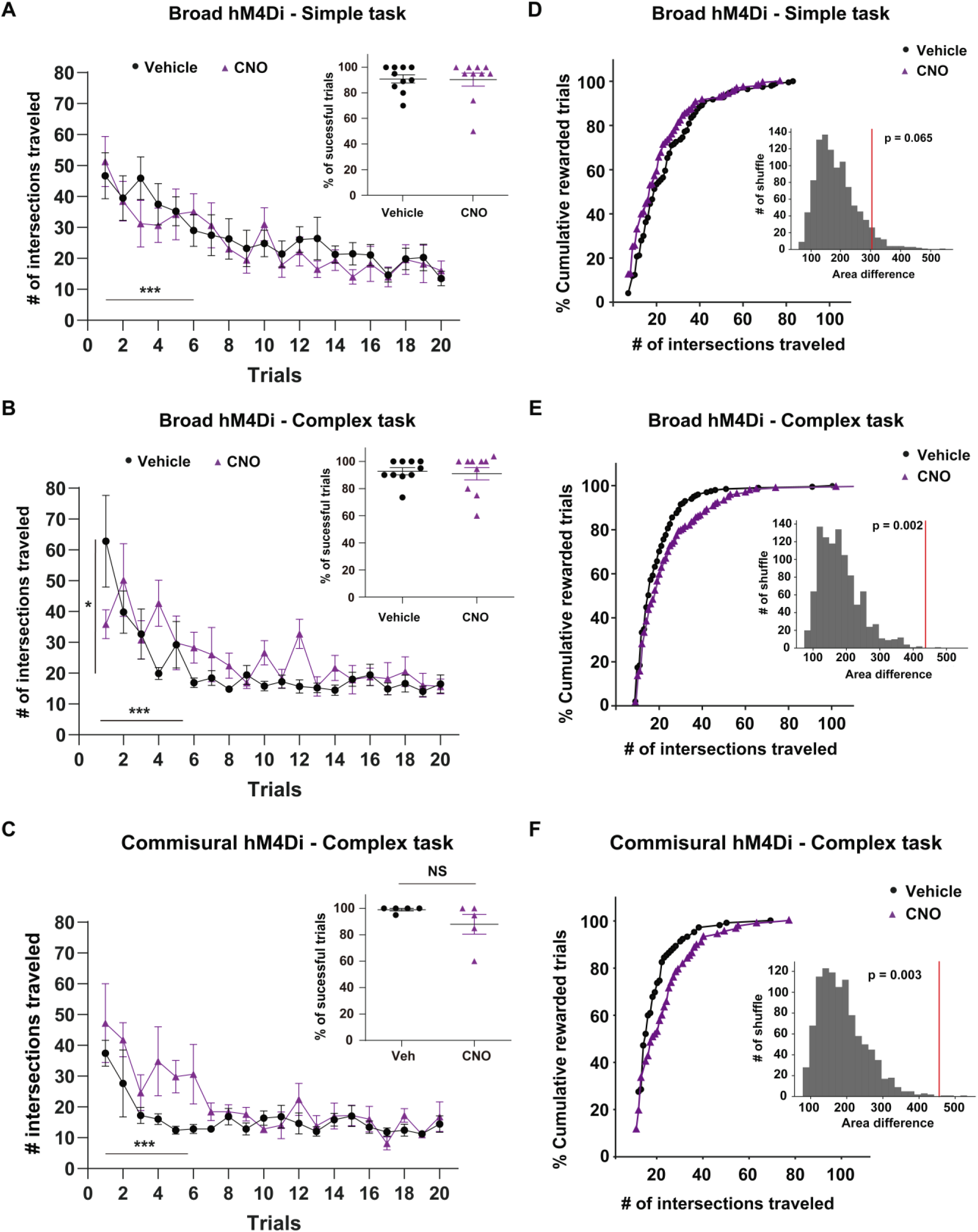
Spatial navigation towards the common novelty in simple and complex tasks. **A**. Vehicle and CNO simple task sessions done for the same 10 mice. The plot showed the number of intersections traveled in 20 trials. Statistical significance for trials and no significant (NS; p = 0.2) for conditions. Inset: Percentage of successful trials for each condition. **B.** Number of intersections traveled over the maze as a function of the 20 trials of the vehicle or CNO complex sessions for the same 10 mice. Statistical significance for trials and conditions. Inset: Percentage of successful trials for each condition. **C.** Number of intersections traveled by the five commissural hM4Di expressing mice along the complex sessions with the vehicle and CNO. Statistical significance for trials and NS (p = 0.11) between conditions. Inset: Percentage of correct trials. NS (p = 0.25) between conditions. **D.** Efficiency of navigation shown by the cumulative percentage of the total of successful trials per condition as a function of the path length done over the maze in the simple task shown in A. N = 173 and 172 total rewarded trials in the vehicle and CNO sessions, respectively. Inset: Tag-shuffles between conditions. The red line is the real area difference between the cumulative distributions with its associated p value (NS in this case). Note that efficiency is not affected by CNO injection in the simple task. **E.** The efficiency plot as panel D but for the same 10 mice in the complex configuration shown in B. N = 181 and 175 total rewarded trials for vehicle and CNO treatments. Note that the CNO distribution was significatively shifted to the right meaning CNO-injected mice did longer paths than in the control condition to reach the same percentage of success. Inset: Distribution of the tag-shuffles differences between the conditions. Significance between the real distributions (red) compared with the differential areas gotten from the shuffles. **F.** Cumulative percentage of the total of correct trials for the commissural hM4Di expressing mice in each session of the complex task. N = 97 and 84 total rewarded trials for vehicle and CNO treatments, respectively. Inset: Shuffles distributions like previous panels. Significance distribution difference between real data (red) compared with the shuffles. Note the same significance deficit in the navigation efficiency when mice were CNO-injected compared with themselves in control condition (vehicle) like it was presented in E.

## RESULTS

### Spatial navigation paradigms in a symmetrical crossword maze

The experimental strategy used to investigate the role of the dentate gyrus in goal-guided behavior within a dynamic environment involved reversible chemogenetic silencing during spatial navigation in a crossword maze. This silencing was achieved by the expression of the synthetic human muscarinic receptor hM4Di, which is exclusively activated by the synthetic agonist clozapine-N-oxide (CNO) (Urban & Roth, 2015). Spatial navigation sessions were monitored in the same mice which, in different goal-guided sessions, received either CNO or vehicle (Fig. 1 A and B). The hM4Di receptor was delivered bilaterally to the dentate gyrus using viral vectors in two ways. A group of mice received an adenoassociated virus (AAV) carrying the hM4Di receptor and mCherry under the CaMKII promoter (AAV-CaMKII-hM4Di-mCherry; N = 10 mice). This AAV labeled most cells in the dentate gyrus and some CA3 pyramidal cells (Fig. 1 C). Another group received a retrovirus expressing the Cre recombinase under the CAG promoter (RV-CAG-Cre) together with an AAV-hSyn-floxed-hM4Di-HA, which expresses hM4Di-HA upon Cre-dependent combination (N = 5 mice). This latter strategy resulted in hM4Di-HA becoming predominantly expressed in commissural fibers of the dentate gyrus (Fig. 1 D, Fig. S 1 A, B).

The crossword maze used in this study was symmetrical to make it equally difficult from any start location. In addition, it was labeled with numbers at each intersection and letters at each path end to generate a coordinate system for subsequent analysis (Fig. 1 E). Thus, the quantity of intersections traveled indicated the animal path length for each trial and the numbers labelled the mouse trajectory on the maze. Each session involved 20 trials, in which mice started alternatively from opposite houses to find a unique goal position. Two navigation challenges were configured. One task was named “simple” and it was the same challenge than previously published (McNamara et al., 2014). The task had fixed barriers along the session, creating two optimal paths of 6 intersections long from opposite houses. Importantly, these paths shared the last streets with the same direction towards the reward position (Fig. 1 F). The other task, called “complex”, was designed as a variant with higher cognitive demand. Hence, there was a mobile barrier that alternated between two positions, each one corresponding to a starting house, generating longer optimal paths with opposite turning directions (Fig. 1 G). In both tasks, the reward position was kept constant along trials within a session. However, the goal place was changed between sessions, constituting a common novelty for each of them.

Adult C57Bl6/J mice were habituated and trained to get familiarized with the goal-guided behavior paradigm in the crossword-like maze used in this study (Fig. 1 A). In the simple task, there were no changes in performance (path length) when comparing the same mice expressing hM4Di injected with vehicle (control) or CNO (inhibited condition). In both control and CNO conditions mice were able to improve their behavior, as shown by the successive decrease in path length and timing along trials (Fig. 2 A and Fig. S. 2 A). CNO-injected mice showed a slight but significant reduction in the number of errors compared to their own performance when injected with vehicle (Fig. S. 2 B). Overall, mice under control or inhibited conditions learn the simple challenge, reaching the reward location in about 90 % of the trials (Fig. 2 A inset). In the complex task, mice could also reach the goal position with a 90 % of success rate in both conditions (Fig. 2 B inset). However, when injected with CNO, mice performed significantly worse in the rest of the analyzed behavioral parameters compared to control conditions (Fig. 2 B and Fig. S. 2 C, D).

To strengthen the notion that the diminished ability to solve the complex task was due to dentate gyurs silencing, mice with restricted hM4Di expression in the commissural fibers were evaluated (Fig. 1 D, Fig. S. 1 A, B). Mice behavior was similar to that observed with broad hM4Di expression since in the vehicle condition they made shorter paths than in the CNO session (Fig. 2 C). In addition, a significant deficit in the timing and number of errors to explore the maze was found during the CNO session compared with the vehicle (Fig. S. 2 E, F). Although specific dentate gyrus inhibition induced a deficit in the animal performance to navigate towards the goal, it did not have any difference in the percentage of correct trials compared with control (Fig. 2 C inset). Therefore, dentate gyrus did not contribute to the animal learning of the simple or the complex tasks, as all mice in both conditions had a significant performance improvement along the trials with the same rate of success. However, this hippocampal region inhibition altered the animal ability to improve their performance with a minimal cost (less time, less travelling, less errors) to finish their trials. These results suggested that the dentate gyrus improved the efficiency in which animals navigate to the reward in the complex task.

In nature, the environments are dynamics so animals must adapt their spatial navigation to achieved their goal to survive. Animals must therefore have evolved to overcome the navigation difficulties posed by their habitats using different strategies with variable levels of efficiency (Nyberg, Duvelle, Barry, & Spiers, 2022). As previously shown in the present study, control mice were faster, displayed fewer errors and traveled shorter distances in the complex configuration than CNO condition (Fig. 2 B – C, Fig. S 2 C - F). Therefore, to quantify these differences, we sought a measure of navigation efficiency. We calculated the cumulative distributions of correct trials as a function of the number of intersections traveled from all mice (Fig. 2 D - F). A shift to the left in the distribution would indicate an efficient behavior, where the success was accomplished at a minimal cost (shortest path to the reward). In the simple task, no difference was found between the control and CNO conditions (Fig. 2 D). In the complex task, the navigation efficiency showed a significant deficit in CNO-treated mice for both hM4Di expression strategies (Fig. 2 E, F). The shift to the right in the CNO curves indicates that mice with a silenced dentate gyrus traveled a longer path to reach the same level of success than the control condition. Therefore, mice with dentate gyrus inhibition become inefficient in the complex task.

The systemic use of CNO was shown to have behavioral effects of its own (MacLaren et al., 2016). Therefore, another group of mice were stereotaxicaly infected with a viral combination expressing only fluorescent markers to test the effects of CNO in the absence of synthetic receptors (N = 6 mice) (Fig. S. 1 C). Under these conditions, the cumulative distributions of the successful trials between vehicle and CNO treatments were very similar in the complex task, highlighting the absence of undesired CNO effects in this study (Fig. S. 1 D, E).

### Dentate gyrus inhibition impairs the efficient redirection to the goal

The navigation mediated by the dentate gyrus was closely investigated focusing in the complex task. In addition, as all mice expressing hM4Di had similar behavioral profiles, the behavioral data of both experimental groups were pooled (Fig. 2 B, C, E and F, Fig. S.2 C - F).

Mice in a labyrinth have previously shown rapid learning, sudden insight and efficient exploration, finding the shortest path to the goal from any point of a maze (Rosenberg, Zhang, Perona, & Meister, 2021). It was also demonstrated that animals displayed this “sudden insight” to quicky know where they are and where to go as a sharp improvement in their navigation performance after their first successful trial in their familiar environment (Grieves, Wood, & Dudchenko, 2016; Tolman, 1948). Given that mice found the reward for the first time in different trials of the session in the present study, the behavior of individual mouse was checked out. After the first correct trial, a vehicle-injected mouse displayed an abrupt decrease in the time searching for the goal in all the subsequent trials (example shown in Fig. 3A). When exposed to CNO, the same mouse improved its navigation time in an alternate pattern along the trials. As the task implied alternating trials of different optimal routes to the same reward place, it became notorious that dentate gyrus inhibition was perturbing the navigation of only one of the routes. Therefore, the spatial navigation for each alternating route in the complex task was analyzed separately for the rest of the work. The route that was solved first was called “match route” and the subsequent alternate path was named “detour route”.

**Figure 3:**
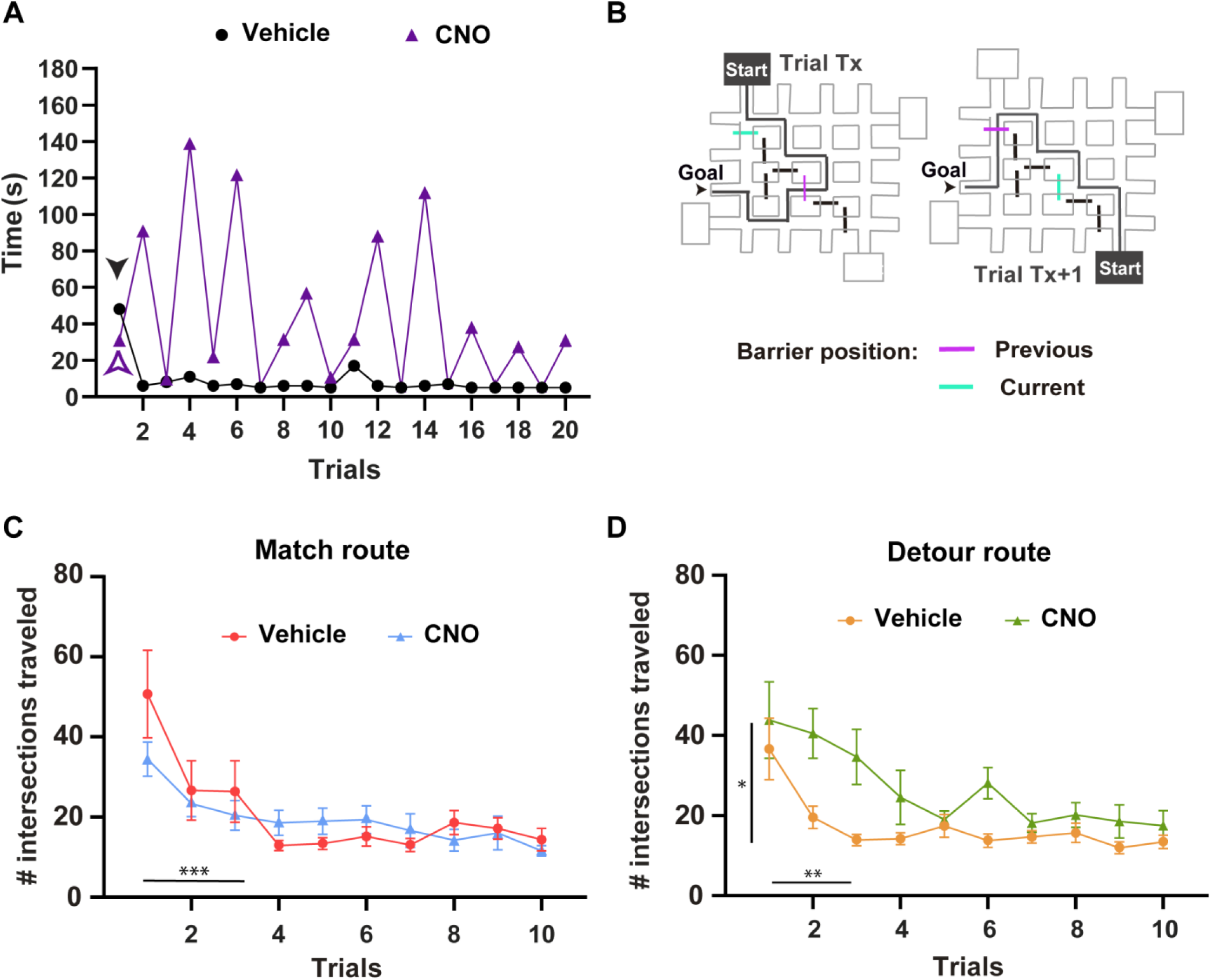
Dentate gyrus support navigation through only one of the alternate routes of the complex task. **A.** Entire spatial goal-guided navigation sessions of one example experimental mouse, injected with CNO for one and with vehicle in the other session. It is shown its navigation timing to finish the 20 trials for each condition. Note this animal found the reward (arrowhead) at the first trial of the vehicle as well as the CNO session. After this, its navigation timing was abruptly reduced and kept low until the end during the vehicle session. However, in the CNO session the same mouse did not have this sudden performance improvement in the following trial from the opposite house. Moreover, its timing showed an alternating pattern along the session. Mostly it was fast for the trials coming out from the same house in which the mouse found the goal (match route) and slow for the alternating pathway (detour route). **B.** Crossword maps examples of a complex task configuration showing that both optimal routes are blocked by the mobile alternating barrier. **C.** Spatial navigation using the match route to find the goal in both conditions represented with the number intersections traveled. Statistical significance for trials and NS (p = 0.24) for conditions. N = 15 hM4Di expressing mice. **D.** Intersections navigated over the maze with the detour route configuration in both conditions. Statistical significance for trials and between vehicle vs CNO, for the same mice and sessions of panel C.

In the beginning of the analysis per route, match and detour trajectories were examined to ensure that they shared the same level of complexity. First, the length of all optimal pairs of alternate routes was assessed (Fig. 1 G). The alternate optimal routes for each navigation session had equal-length path variability for both conditions. They had 10 to 12 intersections from started house to goal place for all pairs (Fig. S. 3 A). Second, the trial number in which mice found the reward for the first time was examined. In general, all mice achieved their goal in the first trial of the session without difference between conditions (Fig. S. 3 B). Lastly, we kept with the experiments where both optimal routes were blocked by the mobile barrier (Fig 3 B). Therefore, in the remaining sections, we will show data from mice that had to change their recently used trajectories to arrive to the same goal place in both paths (11 mice for vehicle, and 13 mice for the CNO condition). While goal-guided navigation over the match route displayed similar performance by vehicle and CNO-injected mice, animals in the CNO condition performed poorly in the first half of the detour trials of the session (Fig. 3 C - D and Fig. S. 3 C - F). This analysis reinforced the concept that the dentate gyrus is not involved in learning the routes, but rather, it improves the navigation quality through an alternative pathway towards the goal location.

The ability to efficiently change the route as a sudden insight was analyzed using the first successful trials of the session aligned to the trial where each mouse found the reward for the first time. Control mice exhibited a significant reduction in the number of errors, the navigation time, and the path length immediately after the first rewarded trial (first match trial vs first detour trial; Fig. 4 A - C). In contrast, the same mice with inhibited dentate gyrus did not show this sudden improvement in the navigation over the first detour trial and neither in the subsequent trials. Therefore, when mice had dentate gyrus inhibition lost their ability to suddenly re direct their spatial navigation to arrive to the same goal location just found (Fig. 4 D - E). Instead of, there was no difference in the navigation quality with or without dentate gyrus in the second correct trial of the match route except for the timing of navigation (Fig. 4 A - C). Thus, there was not a sudden insight in the navigation performance of the match route in any condition. These results suggest that control mice might be using different navigation strategies for each route.

**Figure 4:**
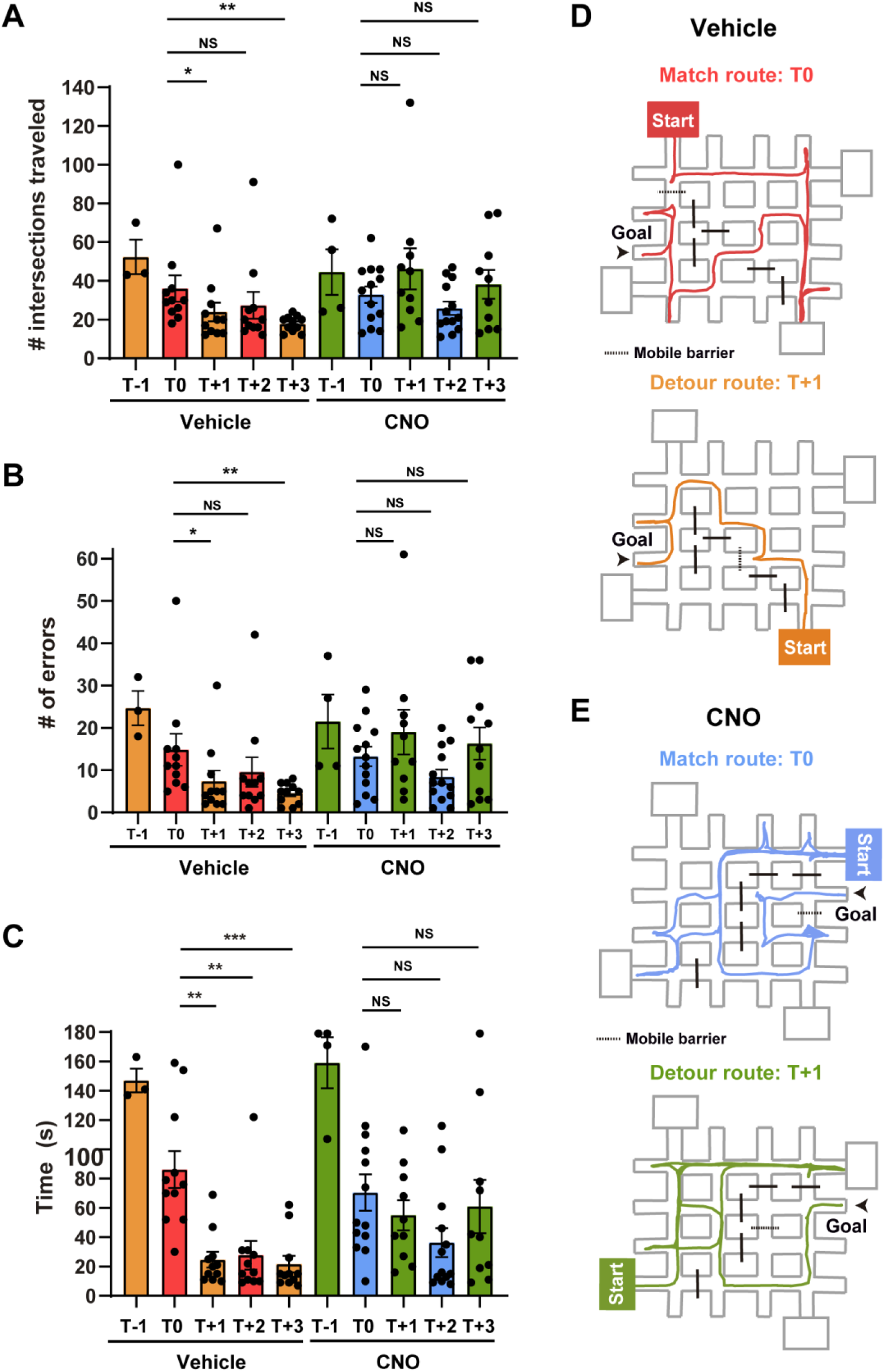
The sudden insight to redirect the spatial navigation through an alternative pathway is dentate gyrus dependent. **A.** Number of intersections traveled on the maze around the first successful trial (T0) during the complex task session in both conditions. Assays are color coded in an alternating pattern to label both optimal routes done to the goal in each condition, from the incorrect trial before (T-1) and the 3 following successful trials (T+1, T+2, T+3) by their chronological order navigated. Note that vehicle-injected mice significantly reduced their path in T+1 as soon as they had found its target location in the previous trial (T0). Instead, the same mice under the CNO effect did not improve their performance in the following trial of the alternating route. Significance was for the following comparisons: T0 vs T+1, T0 vs T+3 and NS for T0 vs T+2 of the vehicle condition and NS for any of the same comparisons for the CNO treatment. N = 11 for vehicle, N = 13 for CNO conditions with identical challenge complex routes. **B.** Peri-event plot as panel A but for the behavioral parameter of number of errors in each condition. The significance and NS were in the same comparisons done for the intersections traveled parameter of panel A. **C.** Navigation timing done in the alternating trials around the event (T0) where mice found the common novelty of the task. Note that vehicle-injected mice significantly improved their timing to navigate to the goal place of both opposite alternating routes as soon as they had found it in T0. In this case there was significance for all subsequent comparisons of successful trials: T0 vs T+1, T0 vs T+2 and T0 vs T+3 in the vehicle condition but still there was NS for all the same comparisons in the CNO condition. Panels A to C show data for the same mice running in the same sessions. **D.** Crossword mazes maps with example trajectories of match (top) versus detour (bottom) routes done by a vehicle-injected mouse in T0 and the following T+1. Note: After the first encounter with the common novelty of the task in T0 the vehicle-injected mouse could travel directly to the just found goal location using a completely different route in T+1. **E.** Routes of goal-guided behavior done by the same example mouse of panel D but CNO-injected. The paths of the same trials shown in panel D; match (top) and detour (bottom) were plotted to highlight one example of the inefficient trajectory done in T+1. In which is notable how the mouse had lost the ability to directly find another way under CNO effect to reach to the same place.

The routing efficiency along the subsequent trials of the session was studied with a cumulative plot of the path length as a function of trials. This analysis was aligned to the first correct trial per mouse, for each route and condition (Fig. 5 A). Remarkably, the navigation efficiency for the match route was similar regardless of dentate gyrus inhibition, from the moment mice found the goal to the end of the session. However, the ability to efficiently arrive to the same reward location through the detour route was strongly impaired when silencing dentate gyrus along all the session. In agreement, a similar effect was observed in the cumulative distributions of all the rewarded trials separated per route and condition, accordingly to the path length (Fig. 5 B). The statistical analysis of these cumulative distributions revealed a significant difference in the routing efficiency between match vs detour when the dentate gyrus was inhibited, while showed no difference for the control condition (Fig. 5 C, D). In addition, the same analysis was consistent with previously observed along the successive trials (Fig. 5 A). There was a strong impairment in navigation quality between conditions along the detour, while no difference was observed in the match route (Fig. 5 E, F). In other words, mice with dentate gyrus manipulation lost their flexibility to efficiently redirect their navigation to a recently found location using a different path in their familiar environment. However, there was no contribution of the dentate gyrus in the first rewarded path, the match route, suggesting that different spatial navigation strategies are supported by different hippocampal regions.

**Figure 5:**
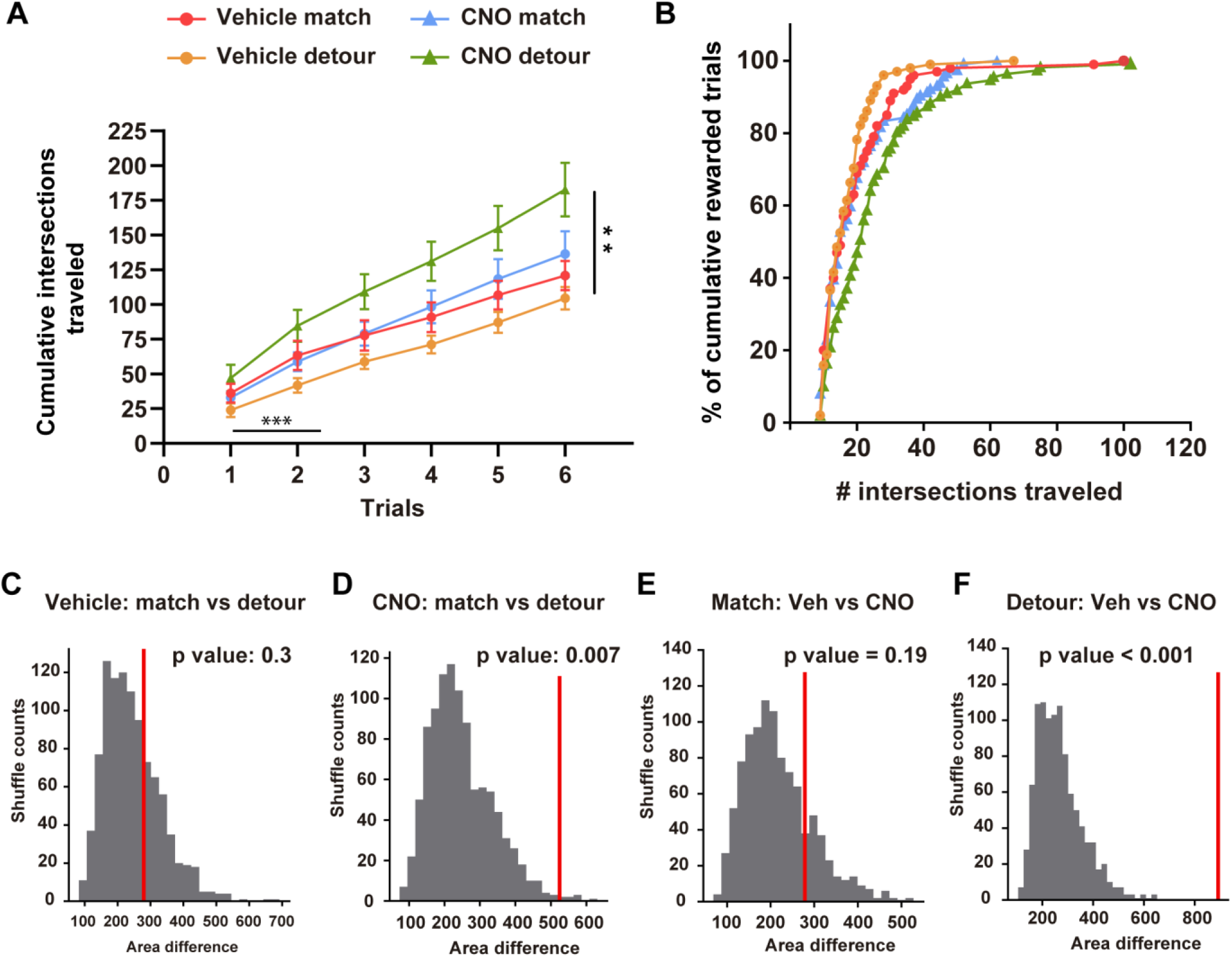
Dentate gyrus provides navigation efficiency in the detour route to the goal. **A.** Cumulative intersections traveled along the successful trials for each condition in the match and detour equal-challenged routes. Rewarded trials for the match trajectory were aligned to the first trial that each mouse found the reward (T0, Fig. 4). Correct assays for the detour path were ordered from the following trial (T+1, Fig. 4) after the first match. Due to this alignment only 6 trials total are presented, because these trials were the ones that had the complete data set of 11 vehicles and 13 CNO-injected mice of equal-balanced routes. Significance for trials and conditions in the detour route. **B.** Efficiency plot of cumulative percentage of the total of successful trials per condition and route as a function of the path length done over the maze. The colored curves have the same code as panel A. Note the CNO distribution of rewarded trials achieved by the detour route is right shifted indicating mice needed to travel more to get the same level of success than controls. N = 100, 100, 94 and 91 total rewarded trials by 11 hM4Di expressing mice in each condition to have balanced distributions of vehicle-matched, vehicle-detoured, CNO-matched and CNO-detoured assays, respectively. To ensure these balanced distributions 2 CNO-injected mice were randomly out of the 13 mice pool. **C.** Tag-shuffles for the succeeded distributions (match vs detour) in the vehicle condition showed NS differences with the real area difference between them (red line with its p associated value) Note: vehicle-rewarded trials had the same efficient distribution in both routes in the complex session. **D.** Distribution of 1000 iterations of tag-shuffle analysis comparing correct trials of match vs detour routes done in the CNO condition resulted in a significant area difference between the real curves (red line, p = 0.007). **E.** Comparison of the area distribution of the shuffles between conditions using the successful assays done in the match route had a NS effect (red line) with the real data difference (red line). **F.** Analysis of the iterations for the tag-shuffles between the distributions of correct trials of each condition using the detour route to the goal presented a robust significance difference (red line). Note: CNO-rewarded trials were significantly inefficient when they were done taking the detour route to the goal.

## DISCUSSION

Spatial learning in the simple task done in the present work was previously shown to be dependent on hippocampal CA1 function (McNamara et al., 2014). The results shown in this study indicate no dependence of this task on dentate gyrus activity, but it supports a requirement for efficient navigation in a more complex version of the task. In the simple as well as the complex version, animals were challenged to find the same goal location using two alternative pathways. However, the pair of optimal pathways of the complex task had completely different path choices from the starting houses to the reward place. Even though this pair of optimal routes displayed similar levels of complexity within one session, silencing the dentate gyrus surprisingly impaired the navigation efficiency only in one of them.

The route that had an inefficient navigation with dentate gyrus inhibition was the detour route of the complex task without any effect on its match pair. To note, it was in this last route where the animals found the reward location for the first time on the maze. Importantly, the goal place was a common novelty between sessions. It has been reported that dopamine release in hippocampal CA1 mediated a memory boost of the experience around the novelty (Duszkiewicz et al., 2019; Yamasaki & Takeuchi, 2017). Therefore, the match route might have been reinforced by dopamine, as it was the first trajectory associated with the common novelty. In contrast, the detour route might have been solved using different cognitive processing supported by the dentate gyrus. In the simple task, the intersections near the goal were common for the match and detour routes. Both routes might have benefited from dopamine release in CA1 as the match route in the complex task. Consistent with our findings, functional dissociations between the dentate gyrus and the CA1 region have already been described (Gilbert et al., 2001).

In the present study, mice with or without dentate gyrus inhibition displayed similar performances in goal-guided navigation of the simple task and in the match route of the complex task. Moreover, mice in both conditions and in both tasks reached the same level of successful trials during the corresponding sessions. These results indicate that the dentate gyrus is not involved in learning the trajectories to the goal. Only mice with a functional dentate gyrus had a sudden spatial insight to reach the goal using a detour route when their recently successful path was blocked. In addition, animals in this control condition had the same level of navigation efficiency in both routes along the session, while it was impaired in the detour route under dentate gyrus inhibition. These dentate gyrus contributions suggest a specific role of this region in cognitive flexibility. This is an adaptative capacity to quickly solve new situations without the requirement of new learning to get the solution. It rather depends on knowledge from previous experiences that had already being acquiree (Nyberg et al., 2022). Consistent with previous work, mice with impaired dentate gyrus function retained their learning capacities but displayed a deficit in cognitive flexibility, required to solve novel situations (Berdugo-Vega, Lee, Garthe, Kempermann, & Calegari, 2021; Burghardt, Park, Hen, & Fenton, 2012; Li et al., 2024).

The mechanism supporting cognitive flexibility is unknown and it might respond to a combination of dentate gyrus capacities. One known ability of the dentate gyrus is the pattern separation process. This function has been shown to be good in detecting small changes over the metric configurations of the cues within the familiar environment. In addition, it has been proposed and evidenced that this region also has the property to associate or bind the objects with their place, an ability called “conjunctive encoding” (Kesner et al., 2015; Lee & Jung, 2017). These capacities together might have generated a detailed map of the relative positions of the local and distal cues of our crossword maze. An updated detailed map for each session of the familiar maze could have been the basis for flexible navigation to reach the goal location, as it has been reported previously (Buzsaki & Moser, 2013; Nyberg et al., 2022). In agreement, previous work has shown that the dentate gyrus was critical for goal-guided spatial navigation based in a map of the environment in a dual solution task (Albrecht et al., 2022).

Another dentate gyrus capacity that could have been combined to provide cognitive flexibility in this study might be the prospective coding ability. This neuronal activity, which anticipates the animals future location, has been shown to be promoted by the dentate gyrus during successful goal-guided navigation (Sasaki et al., 2018). In the complex task of the present study, the mobile barrier blocked the recently successful match route, therefore mice should had predicted a new sequence of intersections to make an efficient detour route to the goal place. Notably, this neuronal activity anticipation has been shown to rapidly adapt to barriers change and predict the correct trajectory to the goal during high-cognitive demand (Dupret, O’Neill, Pleydell-Bouverie, & Csicsvari, 2010; Sasaki et al., 2018; Widloski & Foster, 2022).

In conclusion, the mechanisms for which the dentate gyrus supports cognitive flexibility were not addressed in the present study. However, this region has a lot of cognitive abilities that were discussed in light of the bibliography and in agreement with the results presented here that could be combined in a major adaptative role of this hippocampal area. In this context, the dentate gyrus could have evolved to provide animals with alternative efficient strategies to survive in a changing world.

## ACKNOWLEDGMENTS

We thank members of the A.F.S. and E.K. labs for insightful discussions. PhD Luciana Ferella to perform drug solutions, mice slicing and i.p. injections. PhD María Soledad Espósito to bring us the viral vectors from Silvia Arber laboratory. We also thank the Vivarium technicians and their Director Adriana Fontanals for valuable tips in the animal handling and their lovely daily care of all mice used. V.C.P., E.K., and A.F.S. are researchers in the “Consejo Nacional de Investigaciones Científicas y Técnicas” (CONICET). N.S. and S.B. were supported by CONICET fellowships. This work was supported by grants from Argentine Agency for the Promotion of Science and Technology PICT-2020-0046, PICT-2021-0077 (AFS) and PICT-2016-3611, PICT-2019-00582, Bunge and Born & Fundación Williams Foundations Fellowships (VCP).

