## Supplementary Material for "The dentate gyrus provides flexibility for efficient spatial navigation"

### Supplementary Figure 1

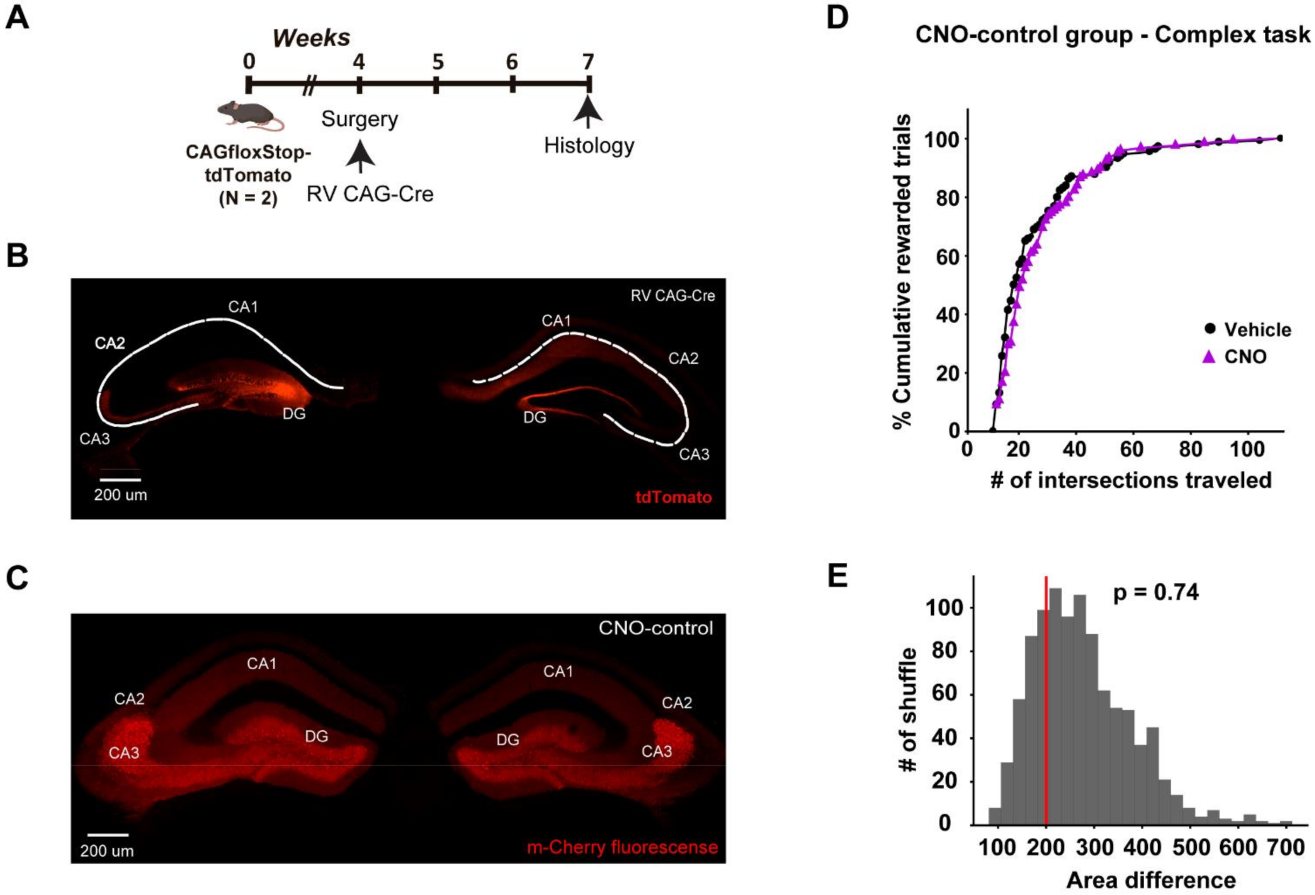


**Supplementary Figure 1: Histological controls and spatial navigation efficiency in CNO-control group. A.** Schematic of the experimental timeline of two CAG-floxStop-tdTomato transgenic mice used to check the RV-CAG-Cre expression. Note the surgery was pointed at the same age as the other experimental animals and 3 weeks of the fluorescent marker expression were allowed. **B.** Representative hippocampal bilateral image of one of the transgenic mice unilaterally infected with the RV-CAG-Cre sliced in a coronal plane. The tdTomato expression was localized in somas, dendrites of dentate neurons in the injected hemisphere (left) and the commissural fibers were labeled in the contralateral dentate gyrus (right). **C.** Histological coronal section showing the bilateral hippocampus of an example CNO-control mouse injected with the adenovirus AAV-hSyn-mCherry in both dentate gyrus region. Note the similar pattern of mCherry expression than there was in the other mice group shown in Fig. 1 C. **D.** Navigation efficiency plot as shown in Fig. 2 D - F, in this case for the CNO-control group in the complex sessions done with vehicle or CNO-injected mice each of them. N = 117 and 108 total rewarded trials in the vehicle and CNO sessions respectively. **E.** Shuffle distribution as done in Fig. 2 D - F. The curves of traveled intersections between conditions showed no area difference between them (red line), meaning that CNO itself is not affecting the animal efficiency.

**Supplementary Figure 2**

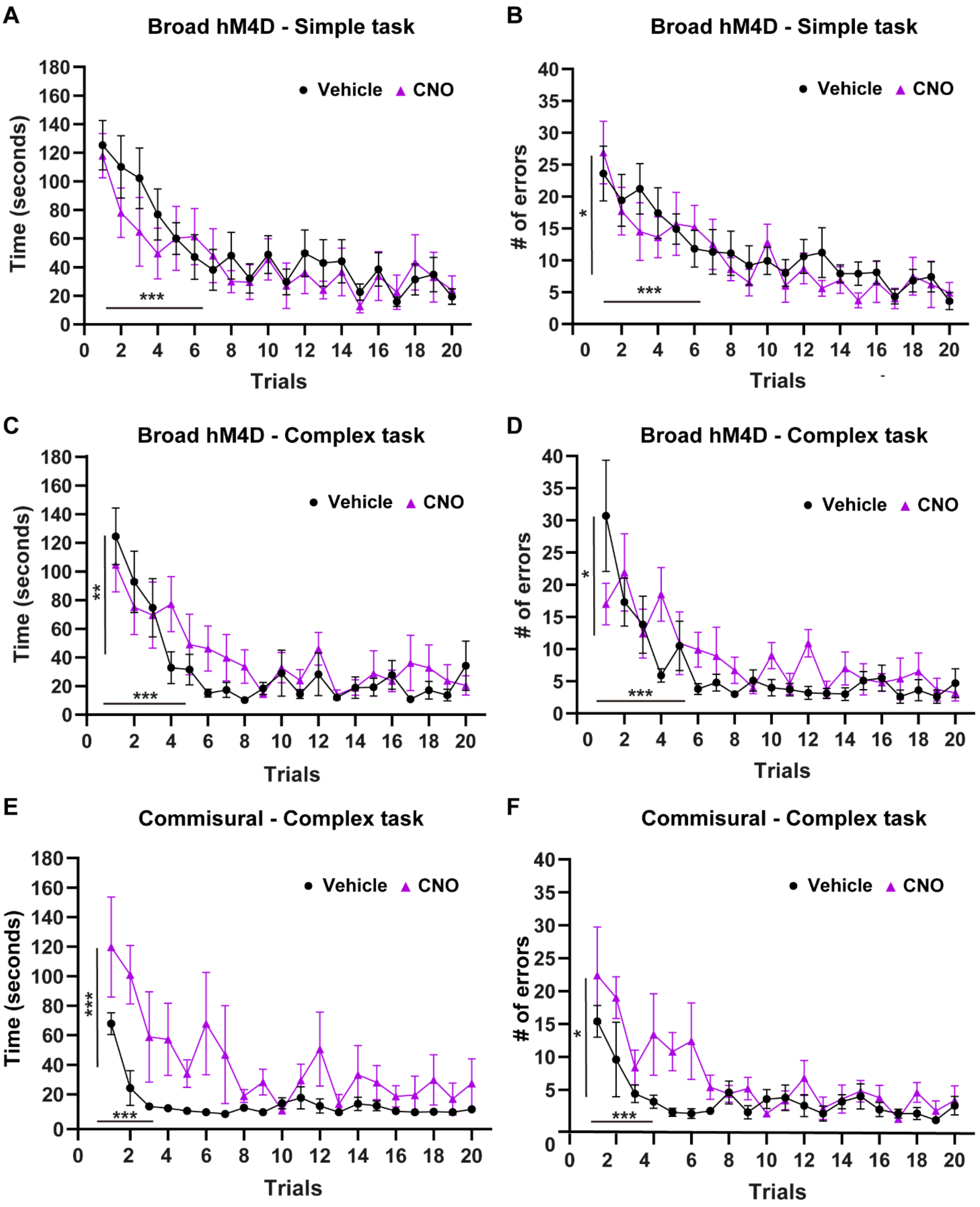

**Supplementary Figure 2: Errors and timing parameters of goal-guided spatial behavior of the experimental groups. A.** Timing of navigation along the 20 trials of the simple task sessions of each condition. Statistical significance for trials and NS (p = 0.15) effect between conditions. **B.** Number of errors in function of the 20 trials of the simple sessions of the same mice shown in A. Statistical significance for trials and for conditions **C.** Timing of navigation over the 20 trials of the complex session of each condition. Statistical significance for trials and for conditions. **D.** Number of errors in both complex sessions (vehicle and CNO). Statistical significance for trials and for conditions. Panel A to D were the same 10 mice as Fig. 2 A, B. **E.** Timing of navigation in function of the 20 trials of each session per condition of the complex task of the dentate gyrus commissural group. Statistical significance for trials and between conditions. **F.** Number of errors along the sessions of the complex task for the same 5 mice as panel E and Fig. 2 C. Statistical significance for trials and conditions.

### Supplementary Figure 3

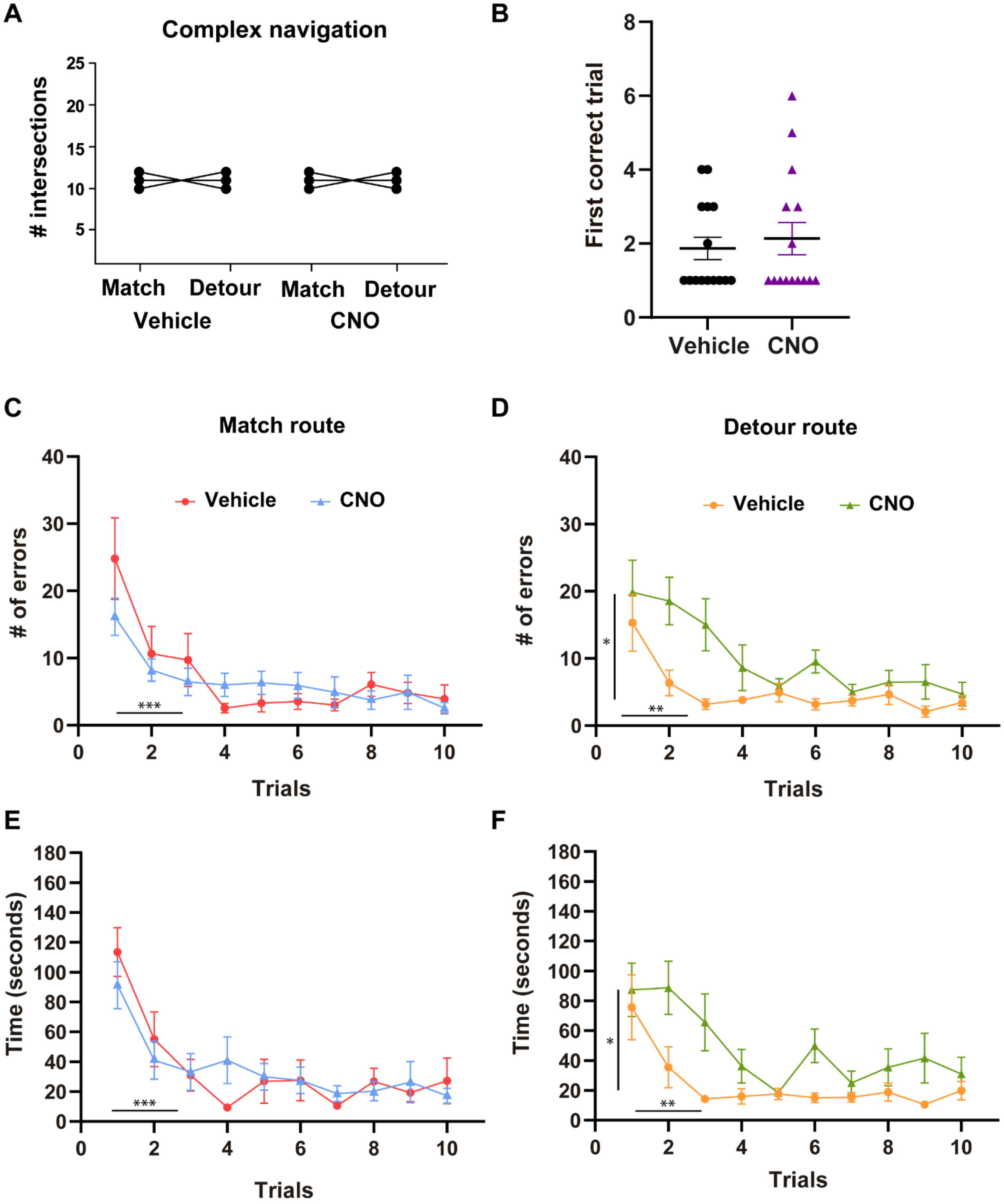

**Supplementary Figure 3: Equal-balanced routes to the same target were differentially impaired with dentate gyrus inhibition. A.** Number of intersections of all the optimal routes from the started house to the goal position used in the complex navigation task of this study. Note that all the routes of the pair had a similar length, 11 ± 1 intersections randomly used for match or detour. **B.** Trial of the session in which each mouse found the reward location novelty for the first time in each condition. Note that most mice found the reward within the first trial of the session in both conditions. N = 15 mice. **C.** Animals mistake using the match route to arrive at their objective in both conditions. Statistical significance for trials and NS ($p = 0.17$) between vehicle vs CNO. N = 11 for vehicle, N = 13 for CNO condition. **D.** Number of errors done looking for the reward position using the detour route along the session of each condition. Statistical significance for trials and conditions. **E.** Timing of navigation over the match route to arrive at the goal location in both conditions. Statistical significance for trials and NS ($p = 0.09$) for conditions. **F.** Duration to finish the trials over the detour route in both conditions. Statistical significance for trials and treatment (vehicle vs CNO). Panels C to F shows data for the same mice running in the same sessions.
